# Controlled Protein-Membrane Interactions Regulate Self-Organization of Min Protein Patterns

**DOI:** 10.1101/2024.03.13.584081

**Authors:** Mergime Hasani, Katharina Esch, Katja Zieske

## Abstract

Self-organizing protein patterns play an essential role in life, governing important cellular processes, such as polarization and division. While the field of protein self-organization has reached a point where basic pattern-forming mechanisms can be reconstituted in vitro using purified proteins, understanding how cells can dynamically switch and modulate these patterns, especially when transiently needed, remains an interesting frontier. Here, we demonstrate the efficient regulation of self-organizing protein patterns through modulation of simple biophysical membrane parameters. Our investigation focusses on the impact of membrane affinity changes on Min protein patterns at lipid membranes composed of E. coli lipids or minimal lipid composition and we present three major results. First, we observed the emergence of a diverse array of pattern phenotypes, ranging from waves to snowflake-like structures. Second, we establish the dependency of these patterns on the density of protein-membrane linkers. Finally, we demonstrate the fine-tuning of snow-flake-like patterns by regulating membrane charge through lipid composition. Our results demonstrate the significant influence of membrane linkage as a straightforward biophysical parameter governing protein pattern formation. Our research points towards a simple yet intriguing mechanism by which cells can adeptly tune and switch protein patterns on the mesoscale.

---

Temporal and spatial control over biochemical patterns is essential for numerous cellular processes, including phenomena such as cell polarization, division, regulation of cell migration directionality, and immunological synapse formation at specific cellular sites. A paradigmatic example of dynamic protein self-organization is the oscillating Min protein system of Escherichia coli (E. coli). In this bacterium, the proteins MinC, MinD and MinE oscillate from pole to pole, creating on time average a concentration gradient with the highest concentration at the cell poles, that inhibits cell division at polar sites^[1–7]^.

The mechanism for Min protein oscillations has been intensively studied. MinD is a membrane binding ATPase^[8,9]^. MinE interacts with MinD and membranes^[10]^ and undergoes a conformational change in a MinD-dependent manner, that results in the release of MinD and MinE from the membrane^[11]^. In vitro reconstitution of Min protein self-organization using purified proteins and model membranes gave rise to the emergence of dynamic wave-like patterns observed through confocal and single molecule microscopy.^[12–15]^ These Min waves have also been shown to inhibit assembly of the cell division protein FtsZ in vitro ^[16–18]^.

Similar dynamic protein patterns, reminiscent of Min protein waves have been observed in eukaryotic cells. Polar actin cytoskeletal patterns in motile cells, for instance resemble segments of self-organizing waves. Experiments in giant cells with increased membrane area support this suggestion by demonstrating dynamic actin cytoskeletal waves on increased membrane areas.^[19]^ Dynamic protein patterns of these eukaryotic cells are often controlled by numerous additional biochemical interactions and therefore harder to reconstitute in vitro. The Min protein system, although not known to require additional regulation, therefore serves as a fundamental model system to explore the emergence of such patterns.

Certain properties of Min proteins itself influence pattern formation, including protein concentrations, genetic modifications and optical on-off control^[20–26]^. The membrane also plays a significant role, affecting Min protein patterns through various properties such as membrane geometry, charge, diffusion, flow, lipid composition, saturation effects and salt concentration ^[15,27–34]^. It has also been shown that the interactions of Min proteins with membranes can affect membrane shape and result in oscillatory beating vesicles^[35–37]^. However how protein linkage to lipid membranes affects pattern formation is less well studied and raises important questions about the robustness of dynamic protein patterns and the hidden potential for controlling the occurrence of protein patterns.

Here we demonstrate a novel approach to tailor Min protein patterns through integration of synthetic membrane interaction linkers. Our approach involved altering the natural interaction between Min proteins and lipid membranes, by purifying Min proteins as fusions with histidine tags and supplementing lipid membranes with DGS-NTA, a component with high affinity for histidine-tags. Our experiments unveil a diverse array of emergent protein patterns, including snowflake-shaped patterns. Furthermore, we explored the impact of membrane charge on these patterns, revealing a fine-tuning effect orchestrated by anionic lipids. Our findings demonstrate the susceptibility of dynamic protein patterns to modulation by fundamental biophysical parameters and provide further evidence that membrane linkage and charge exert significant influence on the regulation of these patterns.

Inspired by the ability of cells to actively regulate interactions between intracellular proteins and lipid membranes, e.g. through posttranslational modifications or expression levels, we sought to investigate whether nature could harness this regulatory capacity to induce controlled variations of mesoscale pattern. To address this question, we focused on biomimetic protein self-organization at lipid membranes and monitored the impact of synthetic membrane linkers on the emergence of protein patterns. We hypothesized that membrane linkage could serve as a fundamental biophysical parameter for modulating mesoscale protein patterns.

To test our hypothesis, we employed the Min protein system, a well-established model system for dynamic pattern formation comprising the proteins MinD and MinE, and generated supported lipid membranes using a vesicle fusion assay (Fig 1A). Although Min proteins in E. coli are not known to be regulated by additional membrane interaction regulators, we chose this protein system due to its archetypical representation of biological pattern and its previous reconstitution on lipid membranes. The supported lipid membrane assay provided a versatile platform for customizing lipid compositions, allowing us to compare protein patterns on membranes assembled from E. coli polar lipids and custom-designed lipid mixtures. E. coli polar lipid extract is a mixture of the lipids phosphatidylethanolamine and the two anionic lipids Phosphatidylglycerol (PG) and cardiolipin and served as a basis for our investigations (Fig. 1). To assess the impact of increased interactions between membranes and Min proteins on protein pattern formation, we supplemented E. coli polar lipids with DGS-NTA, a membrane component which interacts with histidine-tags of recombinant proteins. MinD and MinE were purified as fusion proteins with histidine-tags, labeled using fluorescent dyes and we systematically compared the emergence of Min protein patterns on membranes with and without DGS-NTA (Fig 1B).

**Figure 1.**
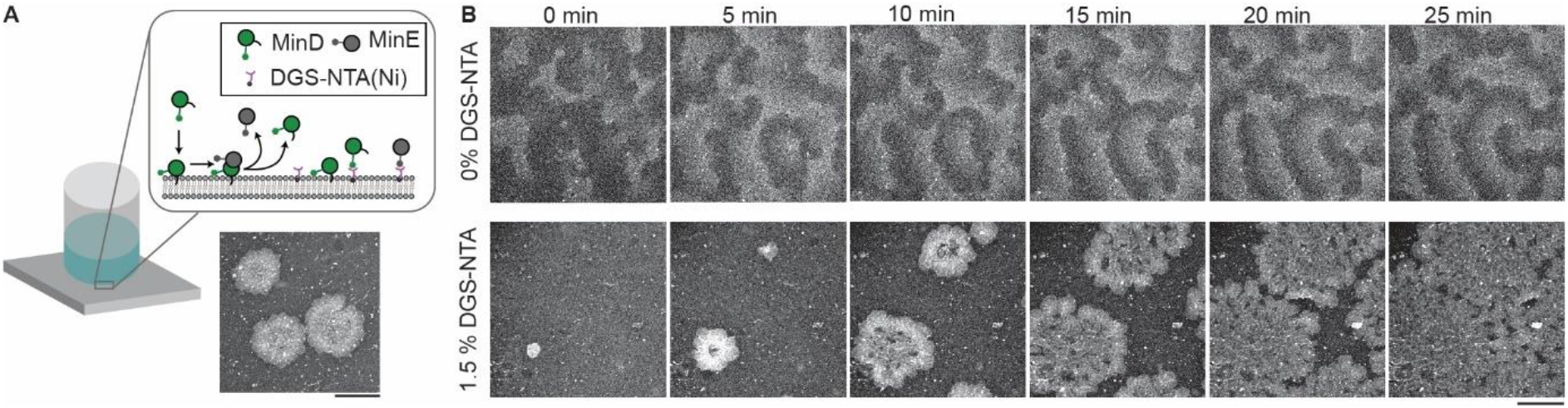
Characterization of Min protein patterns on supported lipid membranes with and without DGS-NTA. A) Schematic of the sample and confocal image of patch-like Min protein patterns in the presence of DGS-NTA. The schematic illustrates a sample chamber and the molecular components of the sample (DGS-NTA, MinD, MinE, lipid membrane). Scale bar: 50 μm. B) Confocal time-lapse images of emerging Min protein patterns on lipid membranes assembled from E. coli polar lipids. Protein concentrations: 1.5 μM MinE doped with 10% MinE-Atto 647, 2.3 μM MinD. Wave-like patterns emerge on membranes without DGS-NTA. On membranes with 1.5% DGS-NTA flower-like patterns emerge and expand until these patterns cover the full membrane surface. Scale bar: 50 μm.

Sequential confocal images of protein pattern formation on these membrane surfaces revealed different patterns in the presence and absence of DGS-NTA (Figure 1B). On membranes without DGS-NTA dynamic protein waves emerged (Figure 1B, upper panel), consistent with previous observations of Min pattern formation on supported lipid membranes.^[13]^ In contrast, the presence of 1.5% DGS-NTA led to the emergence of small protein patches that radially expanded until the whole membrane was covered. Interestingly, these radially expanding protein patches displayed a heterogeneous intensity distribution, giving rise to textures reminiscent of flower blossoms or snowflakes. These results demonstrate the impact of regulating membrane interaction linkers on the modulation of protein patterns. We speculate that cells could achieve such pattern regulation e.g. by modulating the number or activity of membrane proteins that interact with self-organizing protein systems.

Next, we explored the influence of varying concentrations of DGS-NTA on pattern formation and investigated whether snowflake-like patterns could also emerge on minimal membrane compositions. Previous studies demonstrated the emergence of Min protein waves on minimal membranes containing only two lipid species – a neutral and one anionic lipid species^[17]^.

To determine whether snowflake-like patterns could be induced on minimal membranes composed of only two lipid species, namely the neutral lipid 1,2-dioleoyl-sn-glycero-3-phosphocholine (DOPC) and the anionic lipid phosphoglycerol (PG), we supplemented these membranes with varying amounts of DGS-NTA. We found that protein patterns also occur on minimal membranes and that the concentration of DGS-NTA in lipid membranes was a strong determinant for the patterns that emerged (Fig. 2). Membranes lacking DGS-NTA displayed wave-like patterns (Fig. 2A), while membranes supplemented with 1.5% DGS-NTA exhibited snowflake-shaped patterns (Fig. 2C). Intriguingly, membranes with 0.5% DGS-NTA revealed an intermediate pattern phenotype, featuring wave-like segments arranged in flower-like geometries (Fig 2B). Membranes enriched with 3% DGS-NTA displayed protein patches with less-regular shapes (Fig 2D).

**Figure 2.**
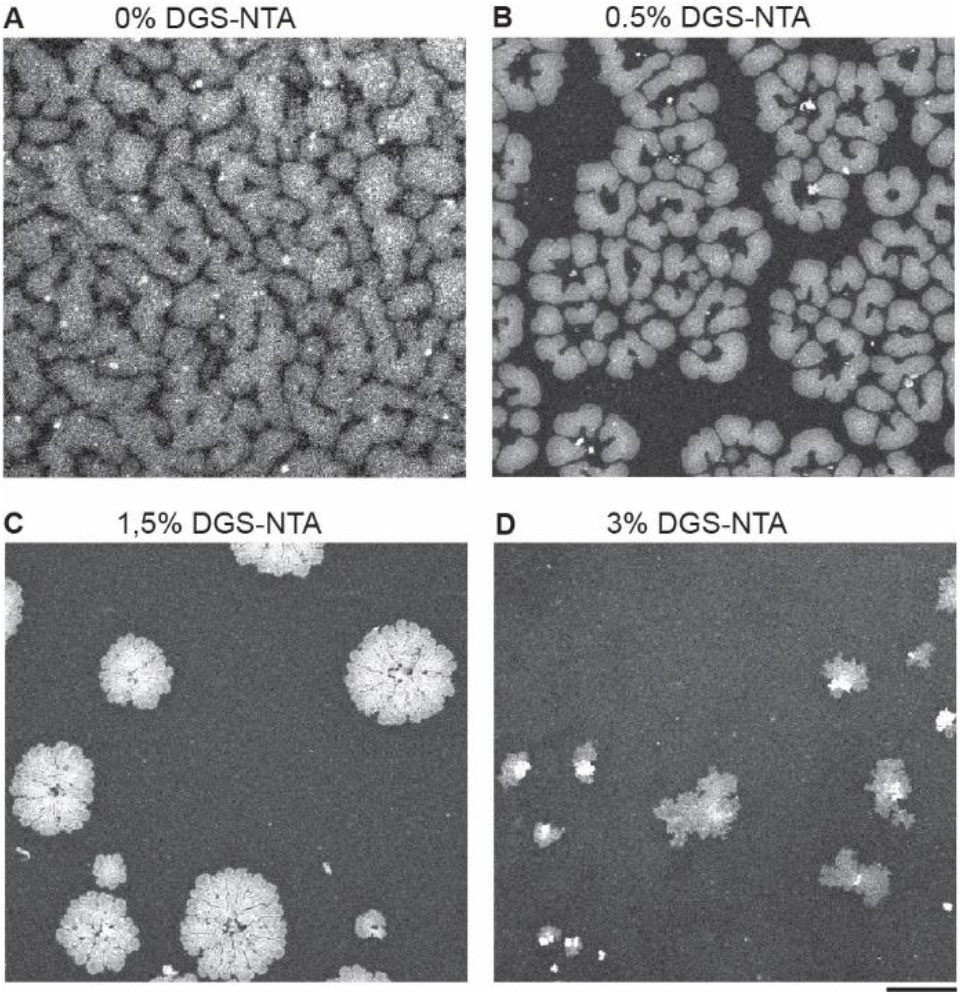
Min protein patterns are modulated by the DGS-NTA concentration in lipid membranes. A-D) Confocal images of Min protein patterns on lipid membrane with minimal lipid compositions (DOPC and E.coli PG at a ratio 1:1, DGS-NTA as indicated in the respective subfigures) 15 min after addition of the proteins. Protein concentrations: 2 μM MinD, 1.5 μM MinE doped with MinE-Atto647. The image contrast was adjusted for each DGS-concentration and is not comparable between subfigures, because protein concentrations were locally much higher when the membrane contained DGS-NTA. Scale bars: 50 μm. A) Wave-like patterns emerge on lipid membranes without DGS-NTA. B) Flower-like patterns emerge on lipid membranes with 0.5% DGS-NTA. C) Snowflake-like patterns emerge on lipid membranes with 1.5% DGS-NTA. D) Irregular patch-like patterns emerge on lipid membranes with 3% DGS-NTA.

To characterize how the expansion dynamics of protein patterns on membranes depend on the concentration of DGS-NTA, we acquired time-lapse confocal images. We found that membranes containing low concentrations of DGS-NTA (0.5%) facilitated the emergence of expanding protein patches, marked by subsequent clearance of proteins in the central area of these patches, resulting in a distinctive wave-fragments-like propagation of patterns (Fig 3A). In contrast, on membranes enriched with high concentrations of DGS-NTA (2.5%), the emerging Min patterns displayed a snowflake-like shape and expanded without clearance of the central area (Fig 3B). Notably, we observed a discernible decrease in the velocity of the pattern fronts with increasing concentrations of DGS-NTA (Fig 3C).

**Figure 3.**
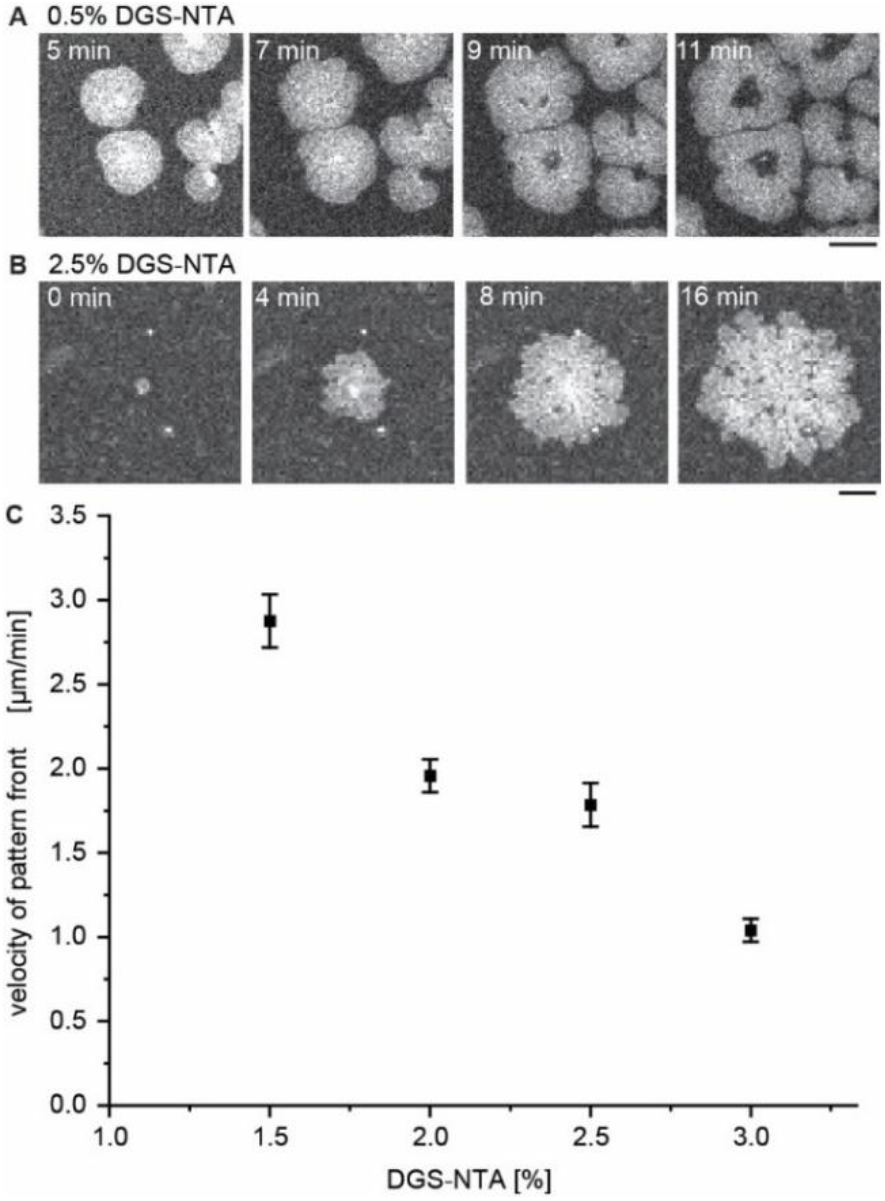
Temporal development of emergent Min protein patterns on membranes with DGS-NTA. A) Confocal time-lapse series of emerging protein structures on a membrane with 0.5% DGS-NTA. Scale bar: 20 μm B) Confocal time-lapse series of an expanding protein patch on a membrane with 2.5% DGS-NTA. Protein patches emerge shortly after adding proteins to the lipid membranes. The patches are initially small and expand over time across the membrane plane. Scale bar: 10 μm. C) The velocity of protein patch boundary was measured for expanding protein on membranes with DGS-NTA concentrations ranging from 1.5% to 3%.

The transition from wave-like patterns over snowflake-shape structures to irregular patch-like features demonstrates the nuanced sensitivity of protein pattern formation to membrane linkers and the capability of membrane interaction linkers to intricately modulate protein self-organization. In addition, these findings unveil a correlation between the concentration of membrane linkers and the expansion speed of protein pattern.

Previous studies demonstrated that negative membrane charge affects the wavelength of Min protein patterns. Therefore, we hypothesized that membrane charge may also modulate the formation of snowflake-shaped Min protein patterns. To test this hypothesis, we devised lipid membranes with 1.5% DGS-NTA and systematically varied the concentrations of the anionic lipid component PG. In the absence of PG, we observed the emergence of nearly spherical protein patches (Fig. 4, 0% PG). However, on membranes containing 30% PG, undulation at the boundary of the protein patches were observed (Fig. 4, 30% PG). Snowflake-shaped geometries emerged on membranes enriched with 50% PG, while those with 70% PG exhibited less regular protein patterns (Fig. 4, 50-70% PG). These results extend the role of membrane charge in shaping protein patterns beyond wave-like patterns to the modulation of snowflake-shaped structures, offering a more nuanced understanding of protein pattern regulation.

**Figure 4.**
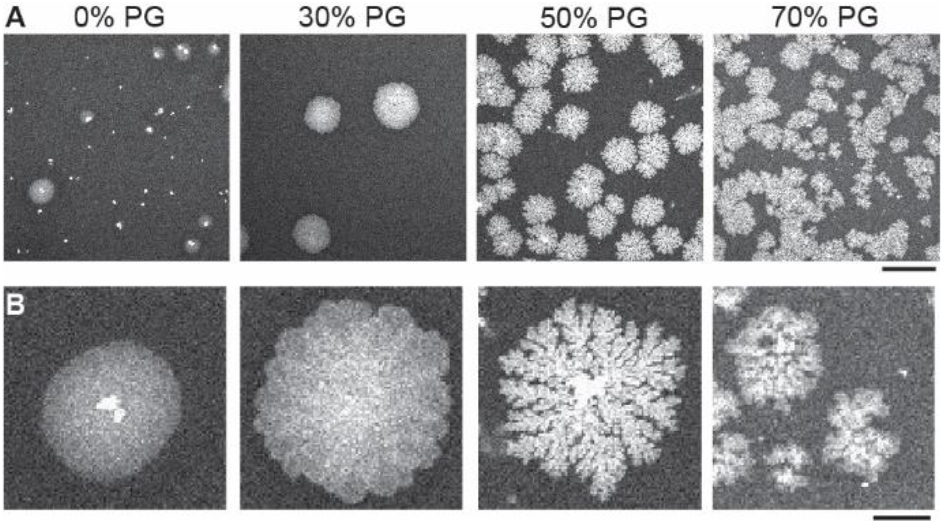
The concentration of anionic lipid in membranes controls the shape of emergent Min protein patterns. A-B) Representative confocal images of Min protein patterns on membranes with 1.5% DGS-NTA and with 0%, 30%, 50% and 70% of the anionic lipid phosphatidylglycerol (PG), respectively. Protein concentrations: 2 μM MinD, 1.5 μM MinE doped with MinE-Atto647. A) Overview images. Scale bar: 100 μm, B) Magnification of protein patterns in A). Scale bar: 20 μm.

In summary, we established an important role of membrane interaction regulation as a highly efficient parameter for modulating various protein pattern phenotypes. Utilizing DGS-NTA as a synthetic membrane anchor to mediate additional interactions between Min proteins and lipid membranes, we demonstrated a spectrum of patterns ranging from waves over snowflakes to patches, dictated by the concentration of DGS-NTA. Furthermore, the manipulation of anionic lipid fractions within DGS-NTA containing lipid membranes allowed for additional fine-tuning of these patterns.

While this study employed the well-established Min protein system as a model to study the pattern-shaping capability of membrane linkers in cell-free environments, the observed pattern modulation by fundamental biophysical parameters suggests broader implications beyond the Min proteins. In eukaryotic systems for instance posttranslational modifications and control over expression levels could serve as plausible regulator for the modulation of mesoscale protein patterns. We believe that our study not only provides insights into mechanisms for nuanced control of protein patterns but also holds promise for biomedical intervention of unwanted biochemical patterns in pathological scenarios.

## Supporting information

Supplemental material: Experimental procedures

## Notes

### Competing Interest Statement

The authors have declared no competing interest.

