## Supplemental material: Experimental procedures for "Controlled Protein-Membrane Interactions Regulate Self-Organization of Min Protein Patterns"

### Table of Contents


1. Experimental Procedures
2. References
3. Author contributions


### 1 Experimental Procedures

**Protein purification and labeling**

MinD was purified with an N-terminal histidine-tag (Addgene # 133621) and MinE was purified with a C-terminal histidine-tag (Addgene #133623) as described previously[1]. In short, the proteins were expressed in *E. coli* strain BL21(DE3). Bacteria were grown at 37°C and induced at $OD_{600}$=0.7 with 1 mM IPTG. The cells were grown for another 3h and harvested by centrifugation. The cells were lysed by sonication and the lysate cleared by centrifugation. The proteins were isolated through Ni-NTA affinity purification. After purification the buffer was exchanged to storage buffer (50 mM HEPES pH 7.25, 150 mM KCl, 10 % glycerol, 0.1 mM EDTA). The storage buffer for MinD was supplemented with 0.2mM ADP) and the proteins were flash frozen in liquid nitrogen. The proteins were stored at -80°C. MinE was labeled with ATTO 647 (Sigma Aldrich). Protein concentrations were estimated with a Nanodrop spectrophotometer.

**Supported lipid membranes**

The reaction chamber for Min protein assays was constructed by affixing a plastic ring onto a class cover slip. For supported lipid membrane formation, small unilamellar vesicles were prepared and fused on the surface of the glass coverslip. To form small unilamellar vesicles lipids (Avanti polar lipids; *E. coli* polar lipid extract or a mixture of DOPC and *E.coli* PG) were mixed with the desired amount of DGS-NTA and 0.05 mol% DiI (Fast DiI, ThermoFisher) in chloroform, dried, resuspended in membrane buffer (25mM Tris-HCl pH 7.5, 150mM KCl) and sonicated at room temperature. Vesicle solution (0.5mg/ml) and 4mM $CaCl_2$ were added to the sample chamber and incubated for 30min at 37°C. The formed membrane was washed with 2ml membrane buffer.

**Min protein imaging**

For reconstituting Min proteins on membrane surfaces, the membrane buffer was exchanged to Min reaction buffer (25mM Tris-HCl pH 7.5, 150mM KCl, 5 mM MgCl2, 2.5mM ATP). The final assay volume was 200µl. Min proteins were added and emerging protein patterns were imaged using a laser scanning confocal microscope (LSM980, Zeiss) with a 20x air objective (Plan-Apochromat 20x/ 0.8, ∞/0.17, Zeiss) at room temperature. ImageJ/Fiji was used for image processing and analysis [2,3].


### 3 Author Contributions

K.Z. designed research, acquired funding and wrote the manuscript. M.H., K.E. und KZ prepared samples, performed research and analyzed the data.
